# Sustained attention and vigilance deficits associated with HIV and a history of methamphetamine dependence

**DOI:** 10.1101/2020.06.03.132522

**Authors:** Nina Pocuca, Jared W. Young, David A. MacQueen, Scott Letendre, Robert K. Heaton, Mark A. Geyer, William Perry, Igor Grant, Arpi Minassian, the Translational Methamphetamine AIDS Research Center (TMARC)

## Abstract

**Background:** Human immunodeficiency virus (HIV)-associated neurocognitive disorders persist in the era of antiretroviral therapy (ART). One factor that is elevated among persons with HIV (PWH) and independently associated with neurocognitive impairment is methamphetamine dependence (METH+). Such dependence may further increase cognitive impairment among PWH, by delaying HIV diagnosis (and thus, ART initiation), which has been posited to account for persistent cognitive impairment among PWH, despite subsequent treatment-related viral load suppression (VLS; ≤50 copies of the virus per milliliter in plasma or cerebrospinal fluid). This study examined the independent and combined (additive versus synergistic) effects of HIV and history of METH+ on the sustained attention and vigilance cognitive domain, while controlling for VLS.

**Methods:** Participants included 205 (median age=44 years; 77% males; HIV-/METH- *n*=67; HIV+/METH - *n*=49; HIV-/METH+ *n*=36; HIV+/METH+ *n*=53) individuals enrolled in the Translational Methamphetamine AIDS Research Center, who completed Conners’ and the 5- Choice continuous performance tests (CPTs).

**Results:** METH+ participants exhibited deficits in sustained attention and vigilance; however, these effects were not significant after excluding participants who had a positive urine toxicology screen for methamphetamine. Controlling for VLS, PWH did not have worse sustained attention and vigilance, but consistently displayed slower reaction times across blocks, relative to HIV-participants. There was no HIV x METH interaction on sustained attention and vigilance.

**Conclusions:** Recent methamphetamine use among METH+ people and detectable viral loads are detrimental to sustained attention and vigilance. These findings highlight the need for prompt diagnosis of HIV and initiation of ART, and METH use interventions.

## 1. Introduction

Modern antiretroviral therapy (ART) has changed human immunodeficiency virus (HIV) from life-threatening illness to chronic disease (Deeks et al., 2013). While ART has also decreased incidence of HIV-associated dementia (Heaton et al., 2011), HIV-associated neurocognitive disorders (HAND) persist (Antinori et al., 2007; Walker & Brown, 2018). Up to half of people with HIV (PWH) present with milder neurocognitive impairment (Heaton et al., 2010). Thus, research should identify how other risk factors prevalent among PWH, including substance use (Dawson-Rose et al., 2017), affect cognitive dysfunction.

Methamphetamine (METH) use and METH dependence (METH+) are prevalent among PWH (Cohen et al., 2016; Dawson-Rose et al., 2017; Hartzler et al., 2017). While amphetamines improve cognitive function among amphetamine-naïve (MacQueen, Minassian, Henry, et al., 2018; Smith & Farah, 2011) and some clinical samples (e.g., attention-deficit/ hyperactivity disorder (ADHD; Baroni & Castellanos, 2015; Sagvolden & Xu, 2008), METH+ has been linked to neurocognitive impairment (Potvin et al., 2018). METH use is also linked to poorer ART adherence among PWH (Moore et al., 2012; Parsons et al., 2013; Passaro et al., 2015), leading to detectable viral loads which are associated with cognitive decline (Heaton et al., 2015). Conversely, viral load suppression (VLS) is associated with stable cognitive function over time (Sanford et al., 2018).

METH+ is also associated with delayed HIV diagnosis and ART initiation (Kuchinad et al., 2016; Passaro et al., 2015), leading to early and more severe immunosuppression. Immunosuppression potentially drives irreversible CNS injury involving frontal systems, resulting in enduring cognitive impairments, that persist despite subsequent VLS (Heaton et al., 2010; Heaton et al., 2011; Muñoz-Moreno et al., 2008). Thus, HIV+/METH+ persons may be at greatest risk of cognitive impairment. Indeed, research points to the combined, deleterious effects of HIV and METH on cognitive function, although it is unclear whether these effects are additive or synergistic in nature (Norman & Basso, 2015; Soontornniyomkij et al., 2016). Differentiating between the additive versus synergistic HIV and METH effects on cognitive function would aid clinicians in selecting appropriate treatments (Brew & McArthur, 2019).

One cognitive function disrupted in PWH and METH+, is sustained attention and vigilance (i.e., the ability to maintain concentration and consistently respond to stimuli; Levine et al., 2008; London et al., 2005; Moran et al., 2014; Moran et al., 2019; Morgan et al., 2014; Potvin et al., 2018). Sustained attention and vigilance underlie other cognitive functions (e.g., learning; Fortenbaugh et al., 2017) and implicated in ART adherence (Hinkin et al., 2002). Research has yet to determine how METH+ affects sustained attention and vigilance in PWH. Using Conners’ continuous performance test (CPT; Conners et al., 2000), Levine et al. (2006) found that cocaine and amphetamine (stimulants that share pharmacological similarities with METH) are associated with poorer sustained attention and vigilance, among PWH. The absence of an HIV-negative (HIV-) group in Levine et al. (2006) precludes inferences regarding the additive versus synergistic effects of HIV and stimulants on sustained attention and vigilance. Further, although a valid and reliable measure (Egeland & Kovalik-Gran, 2010), Conners’ CPT has not been adapted for use in non-humans, precluding the utilization of preclinical studies to determine mechanistic underpinnings of observed deficits (Young et al., 2009). Such studies are difficult to conduct with human participants (Markou et al., 2009).

Ultimately, research is needed to determine the nature of HIV and METH effects on sustained attention (Brew & McArthur, 2019), using a validated, cross-species measure such as the 5-choice CPT (5C-CPT; Barnes et al., 2012; MacQueen, Minassian, Kenton et al., 2018; Young et al., 2009; Young et al., 2013; Young et al., 2017; Young et al., 2019; Young et al., 2020). Thus, this study evaluated the additive versus synergistic effects of HIV and METH on sustained attention and vigilance, controlling for VLS. It was hypothesized that HIV and METH would have a synergistic deleterious effect on sustained attention and vigilance in PWH.

## 2. Material and methods

### 2.1. Participants

Participants were drawn from the Translational Methamphetamine AIDS Research Center (TMARC), an interdisciplinary research program examining HIV and METH+ effects on neurocognitive function. This study used data from 205 participants who completed Conners’ and 5C-CPT (18-87 years, median age=44; see Table 1 for participant characteristics). Most PWH were male (*n*=97, 95%), consistent with the gender distribution of HIV infection in California (California Department of Public Health, 2019).

### 2.2. Procedures

Participants were recruited from the community, HIV clinics, and substance use treatment programs in San Diego if they: (1) were aged ≥18 years; (2) met DSM-IV criteria for METH+: (i) in the past 18 months, or (ii) in their lifetime, and met one DSM-IV abuse criterion in the past 18 months; or (3) had no history of METH use disorder. Participants with a head injury with loss of consciousness >30 minutes, or a medical, serious psychiatric or neurological condition not associated with HIV and linked to neurocognitive deficits (e.g., Hepatitis C infection), or who met past 12-month DSM-IV dependence criteria for another substance (excluding nicotine), were excluded.

Participants provided informed consent and were assessed between 2014 and 2019. Participants provided demographic and substance use information. The Composite International Diagnostic Interview (CIDI) assessed substance use disorders, while the Diagnostic Interview Schedule for DSM-IV (DIS) assessed mental health disorders. Participants underwent a blood test to confirm HIV status and examine plasma viral loads. ART use and nadir CD4 count information were also collected. Participants were not provided with explicit instructions to abstain from substances, however, they completed a urine toxicology screen prior to cognitive testing, which included 5C-CPT and Conners’ CPT. Participants were reimbursed for their time. Procedures were approved by the university Institutional Review Board.

### 2.3. Cognitive Tasks

Cognitive tasks were presented on a 56cm CRT Dell PC computer screen (60cm from participant), using E-Prime2 software (Psychology Software Tools, 2012) for stimulus presentation and data acquisition.

#### 2.3.1. 5C-CPT

Participants responded using an arcade joystick that returned to center after each response. Trials consisted of target (single circle) or non-target (five circles) stimuli that appeared for 100ms, behind and arc of 5 lines. Participants were able to respond for ≤1s after the stimuli disappeared (limited hold). Trials were separated by 0.5, 1, or 1.5s inter-trial interval (ITI), programmed in a quasi-random manner so the same ITI did not appear in more than 3 consecutive trials. Participants completed 12 practice trials (10 target, 2 non-target) to demonstrate understanding of task instructions, before completing the task (225 target and 45 non-target trials).

Responding to target stimuli in the indicated direction was recorded as a hit. Inhibiting responses to non-target stimuli was a correct rejection. Failure to respond to a target was an omission, while responding to a location other than the circle was registered as incorrect. Failure to inhibit responding during non-target trials resulted in a false alarm (FA). Responding before stimuli appeared was a premature response (PR). Outcomes were calculated from these measures using signal detection theory based on hit rates (HR), FA rates (FAR), accordingly:

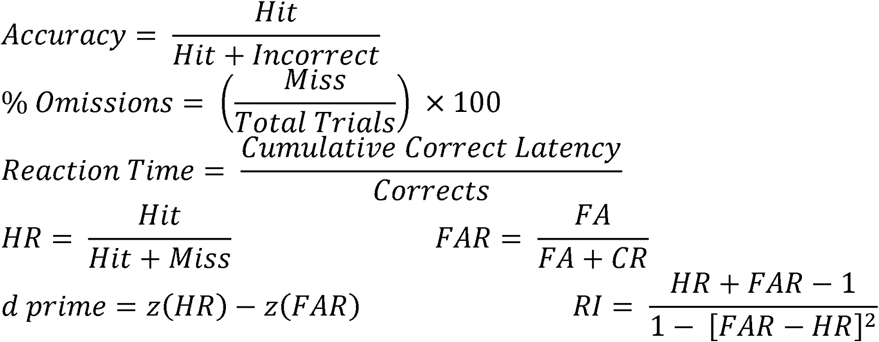

Responsivity index (RI) measured ‘tendency to respond’, where low numbers indicated a conservative and high numbers indicated a liberal response strategy (Frey & Colliver, 1973; Sahgal, 1987). d prime (primary outcome) is a standardized measure (denoted by “z”) of sensitivity to appropriate responding. Accuracy (moving the joystick in the direction of the target), RI, HR, omissions, FA, reaction time (RT), variability in RT, and PR were examined as underlying drivers for performance (secondary outcomes).

#### 2.3.2. Conners’ CPT-2

Participants were presented with letters (separated by an Inter-Stimulus Interval of 1, 2, or 4 seconds) and instructed to use a keyboard to respond to all letters except “X”, where they were instructed to withhold responses. Participants completed 18 blocks of 20 trials (total of 360 trials). The non-target stimulus (“X”) appeared on 10% of trials. Target trials were a hit (response) or omission (no response) and non-target trials were a false alarm (response) or correct rejection (non-response). Response latencies were recorded. d prime measured sensitivity to appropriate responding (primary outcome), while FA, omissions, and RT were examined as drivers of performance (secondary outcomes). ADHD confidence index, a composite measure that prospectively predicts ADHD diagnosis (Breaux et al., 2016), was also examined.

### 2.4. Statistical Analyses

Demographic, mental health, and substance use differences across HIV and METH groups, ART use, nadir CD4 count, and VLS across PWH, and convergence between the 5C-CPT and Conners’ CPT on d prime, omissions, and false alarms were evaluated via regressions. Concurrent main (i.e., additive) and interactive (i.e., synergistic) effects of HIV and METH on overall sustained attention (d prime) and drivers of performance (e.g., omissions, false alarms), were examined via separate forced entry regressions. Drivers of performance were examined regardless of d prime effects. Significant interactions were followed-up via ANOVA to examine group differences. Mixed effects linear regressions examined main and interactive effects of HIV and METH on vigilance, with performance split across three (5C-CPT) or six (Conners’ CPT) blocks. A random intercept accounted for the repeated measures nature of the data, with trial block, HIV, and METH modelled as fixed effects. Model fit was evaluated via −2 log likelihood changes.

Continuous predictors were z-standardized and bootstrapping corrected for deviations from normality. Regressions controlled for age, sex, education, and VLS, given their effects on cognitive function (Heaton et al., 2015; Maki et al., 2018; Samson & Barnes, 2013; Satz et al., 2011). A Bonferroni alpha correction for the greatest number of comparisons within a measure (i.e., eight comparisons in the secondary outcomes for the 5C-CPT, α = .05/8 = 0.006), was applied to correct for family-wise error rate. No alpha correction was applied to d prime analyses, given that it was the primary outcome. Data were analyzed using SPSS (v26.0).

## 3. Results

### 3.1. Data screening and group differences

Group differences are reported in Table 1. There were no significant HIVxMETH interactions. METH+ subjects were younger (*B*=-3.974, *SE*=1.846, 95%CI[-7.583, −0.364]) and less educated (*B*=-1.257, *SE*=0.345, 95%CI[-1.922, −0.573]), than METH-subjects. PWH were more likely to be male (OR=0.075, 95%CI[0.015, 0.183]), than female. HIV+/METH+ subjects did not differ from HIV+/METH-subjects on ART use (OR=0.747, 95%CI[0.221, 2.529]), or nadir CD4 count (*B*=-45.552, 43.726, 95%CI[-127.586, 43.560]); however, HIV+/METH+ were less likely to have VLS in plasma (i.e.,≤50 copies of the virus per milliliter; OR=0.298, 95%CI[0.075, 0.768]). HIV+/METH+ subjects did not differ from HIV-/METH+ subjects on any methamphetamine use characteristics. 5C-CPT d prime, omissions, and false alarms were positively associated with Conners’ CPT d prime (*B*=0.062, *SE*=0.029, 95%CI[0.003, 0.113]), omissions (*B*=8.891, *SE*=2.046, 95%CI[4.981, 12.601]), and false alarms (*B*=1.404, *SE*=3.805, 95%CI[1.062, 16.352]), respectively. Two participants (<1%) had missing data on VLS and were excluded from analyses, resulting in a final analytical sample of *N*=203. 16 METH+ participants had a positive urine toxicology screen for methamphetamine (UTOX(meth)+). Thus, analyses were also run without UTOX(meth)+ participants.

### 3.2. Overall effects of HIV and METH on the 5C-CPT

There were no HIV effects on any outcome (Table 2 and Figure 1). METH+ was associated with lower d prime. There were no other METH or HIVxMETH effects.

#### 3.2.1. 5C-CPT Analyses excluding participants with a positive urine toxicology screen for methamphetamine

There were no significant main or interactive effects of METH or HIV on any outcomes (Supplemental Table 1).

### 3.3. Effects of HIV and METH, aggregated across trial blocks on the 5C-CPT

The main effects model (Table 3 and Figure 2) had better fit, relative to a two-way interaction model (Table 3), thus the three-way interaction model was not examined. There was a significant effect of trial block on d prime, reflecting a vigilance decrement, driven by reduced HR, and higher omissions. Responses also slowed across trials. There were no other significant block effects. VLS was associated with higher d prime and more premature responses. Controlling for VLS, PWH performed comparably on vigilance (d prime), but exhibited less accuracy and slower RT, compared to HIV-participants. There were no other HIV effects. METH+ was associated with worse vigilance (d prime), driven by lower HR and more omissions. METH was not associated with accuracy or RT.

#### 3.3.1. 5C-CPT by block analyses excluding participants with a positive urine toxicology screen for methamphetamine

The main effects model (Supplemental Table 3) had better fit, relative to a two-way interaction model (Supplemental Table 2). METH+ was not significantly associated with any outcomes, while PWH had slower reaction time, compared to HIV-participants. HIV was not significantly associated with accuracy after removing UTOX(meth)+ participants.

### 3.4. Overall effects of HIV and METH on Conners’ CPT

There were no significant effects of HIV or METH on any outcomes (see Table 5 and Figure 3). There was a significant HIVxMETH interaction on ADHD confidence index (prior to Bonferroni adjustment), whereby HIV-/METH+ (*M*=67.750, *SE*=3.205) participants had a higher ADHD confidence index than HIV-/METH- (*M*=54.142, *SE*=2.335) participants (*Mean*^diff^=13.608, *SE*=3.879, 95%CI[6.044, 22.167]). There was no significant difference between HIV+/METH+ (M=62.938, SE=2.780) and HIV+/METH-(M=59.939, SE=2.647) on ADHD confidence index (Mean^diff^=2.998, SE=3.555, 95%CI[-4.410, 9.749]). There were no other significant HIVxMETH effects.

#### 3.4.1. Conners’ CPT Analyses excluding participants with a positive urine toxicology screen for methamphetamine

There were no significant main effects of METH+ or HIV on any outcome variables. There was a significant HIVxMETH interaction on ADHD confidence index (prior to Bonferroni adjustment). HIV-/METH+ (*M*=65.472, *SE*=1.309) participants had a higher ADHD confidence index than HIV-/METH- (*M*=54.459, *SE*=0.899) participants (Mean^diff^=11.013, *SE*=1.693, 95%CI[7.634, 14.340]). There was no significant difference between HIV+/METH+ (*M*=59.390, *SE*=1.198) and HIV+/METH- (*M*=60.575, *SE*=1.034) on ADHD confidence index (Mean^diff^=-1.185, *SE*=1.425, 95%CI[-3.945, 1.681]; Supplemental Table 4).

### 3.5. Effects of HIV and METH, aggregated across trial blocks on Conners’ CPT

The main effects model (Table 7 and Figure 4) had better fit, relative to a two-way interaction model (see Table 6), thus the three-way interaction model was not examined. Omissions, FAs, and RT error significantly increased across blocks, while RT remained stable. VLS was associated with less omissions. Controlling for VLS, HIV was only associated with slower RT. METH+ participants had more omissions, RT error, and slower reaction time, relative to METH-participants. METH+ was not associated with FA.

#### 3.5.1. Conners’ CPT by block analyses excluding participants with a positive urine toxicology screen for methamphetamine

The main effects model (Supplemental Table 6) had better fit than a two-way interaction model (Supplemental Table 5). METH+ was not significantly associated with any outcomes. HIV+ was only associated with slower RT.

## 4. Discussion

This study examined the additive versus synergistic effects of HIV and METH on sustained attention and vigilance. METH+, but not HIV, was associated with sustained attention and vigilance deficits, after controlling for VLS, suggesting a lack of additive HIV effect. There was no synergistic effect of HIV and METH. METH+ was associated with poorer sustained attention and vigilance, driven by lower hit rate (5C-CPT) and more target omissions (5C-CPT and Conners’ CPT). METH+ was also associated with slower reaction time and greater reaction time error (Conners’ CPT). The main effects of METH+ on sustained attention and vigilance disappeared after removing UTOX(meth)+ participants, highlighting the role of recent METH in this relationship.

Contrary to hypotheses, PWH did not have worse sustained attention and vigilance relative to HIV-participants, displaying only a decrement in accuracy, which disappeared after removing UTOX(meth)+ participants. PWH displayed reaction time slowing (both CPTs), which persisted after removing UTOX(meth)+ participants. Contrary to hypotheses, there was no HIVxMETH+ interaction on sustained attention and vigilance. However, there was a significant HIVxMETH interaction on ADHD confidence index. HIV-/METH+ participants had a higher ADHD confidence index, relative to HIV-/METH- participants (prior to Bonferroni correction), even after removing UTOX(meth)+ participants.

Recent METH use among METH+ people, rather than METH+ alone, was associated with sustained attention and vigilance deficits. This result extends Levine et al. (2006), highlighting sustained attention and vigilance deficits with recent stimulant use, irrespective of HIV. Results also extend Basterfield et al. (2019), who found rebound effects for certain cognitive domains among recently abstinent METH+ people. Further research should examine what aspect of recent use (acute intoxication, dosage, residual effects, or withdrawal) is associated with impairment and the direction of this relationship. Namely, whether METH+ people with greater sustained attention and vigilance deficits are likely to have used METH recently (e.g., as an attempt to self-medicate attentional or other problems), or whether recent METH use among METH+ people confers sustained attention and vigilance deficits. These studies are important given that amphetamine administration in healthy participants improves 5C-CPT performance in humans, mice (MacQueen, Minassian, Kenton et al, 2018), and rats (Young et al, 2020). Chronicity of METH use may obviate the benefits of acute effects.

Although the lack of METH+ effects after removing UTOX(meth)+ participants are at odds with Potvin et al., (2018), it should be noted that the studies included in that meta-analysis differed on participant METH use recency. Some studies required participants to be UTOX(meth)+ during cognitive testing, whereas others included recently abstinent METH+ participants. Since Potvin et al., (2018) did not control for METH use recency, it is not possible to disentangle whether METH+ alone, or recent METH use among METH+ people (as found in this study), affected cognitive function.

Several reasons may account for the lack of synergistic HIV and METH effect on sustained attention and vigilance. METH+ may not have necessarily pre-dated HIV seroconversion. Thus, HIV diagnosis (and ART initiation) may not have been delayed, sparing this group from enduring brain injury and cognitive impairment. This notion is supported by Montoya et al. (2016) who found 20% of HIV+/METH+ persons initiated METH use following HIV diagnosis. Alternatively, the lack of synergistic HIVxMETH effects may indicate that HIV+/METH+ are no more susceptible than HIV+/METH- to early immunosuppression (leading to enduring cognitive impairment). Indeed, our results found no difference in nadir CD4 count between the HIV+/METH- and HIV+/METH+ groups. However, it is unclear whether these similar nadir CD4 counts are attributable to HIV diagnosis predating METH+ onset, or whether HIV+/METH+ persons are truly no more susceptible to early immunosuppression, than HIV+/METH-individuals. Further research is required.

Nonetheless, there was a HIVxMETH interaction on Conners’ CPT. HIV-/METH+ participants had a higher ADHD confidence index, than HIV-/METH- participants, a group difference not found using the DIS for DSM-IV, which relies on self-report of symptoms. This discrepancy may underscore the sensitivity of Conners’ CPT ADHD confidence index in capturing the elevated rates of ADHD among METH users, noted in previous studies (Bordoloi, Chandrashekar, & Yarasi, 2019) and observed in our HIV-/METH+ group. Further research should aim to replicate these results, which were not significant following Bonferroni correction and thus, should be interpreted with caution.

HIV was only consistently associated with psychomotor slowing. This finding may indicate that while an impairment in reaction time is still detectable, the current sample of PWH had lower levels of cognitive impairment, than found in previous, virally suppressed PWH (Sanford et al., 2018). The discrepancy in findings may be attributable to the generally high nadir CD4 counts across both HIV+ groups in the current study, reflecting on average, the absence of previous immunosuppression. Alternatively, these results may be attributable to the fact that while untreated HIV has an irreversible, detrimental impact on psychomotor and some cognitive functions (Muñoz-Moreno et al., 2008), sustained attention and vigilance may be spared from enduring deficits. Current results may also indicate that while ART has pro-cognitive effects on some cognitive domains, it has little effect on psychomotor function, the most prominent deficit associated with advanced HIV (i.e., acquired immunodeficiency syndrome; Reger et al., 2002). Both notions are supported by Eggers et al. (2017) who posited psychomotor slowing to be a salient feature of HAND, despite VLS. Alternatively, psychomotor slowing may be indicative of a speed/ accuracy trade-off, whereby PWH compensate for cognitive impairment by slowing responding to prioritize accuracy, as found previously (Kronemer et al., 2017). Animal models are well-positioned to further test these hypotheses.

### 4.1. Implications

Current and previous findings highlight the importance of VLS in cognitive function among PWH (Heaton et al., 2015), reiterating the need for prompt HIV diagnosis and ART initiation (and maintenance). Routine opt-out screening may be a cost-effective means of increasing prompt HIV diagnosis (Krueger et al., 2019; Sanders et al., 2005). Peer-to-peer social support increases ART adherence among substance-using and non-using PWH (Horvath et al., 2013; Hosek et al., 2018; Kerrigan et al., 2018). Given that sustained attention and vigilance deficits were less pronounced in recently abstinent METH+, clinicians should treat METH+ to halt cognitive impairment associated with recurrent use. Cognitive behavioral therapy and contingency management have some efficacy in treating METH+ (Lee & Rawson, 2008; Rawson et al., 2006; Roll, 2007). Such suggestions, while not novel, are strongly supported by the current data.

Current results have implications for the assessment of sustained attention and vigilance. Significant associations were observed between the 5-Choice and Conners’ CPTs. Both CPTs captured vigilance deficits seen in METH+ persons and slowed reaction time seen in PWH. These findings reiterate the validity and provide further psychometric support for 5C-CPT as a measure of sustained attention and vigilance among PWH and METH+ persons. Using 5C-CPT allows for comparisons with future animal research aimed at delineating mechanisms underpinning the deficits seen in this study.

### 4.2. Strengths and limitations

We controlled for age, sex, education, and VLS, which are associated with cognitive function (Heaton et al., 2015; Maki et al., 2018; Samson & Barnes, 2013; Satz et al., 2011). We excluded people with current substance dependence (excluding nicotine), a history of head injuries, medical, psychotic, or neurological conditions known to impact cognitive function (Cunha et al., 2013; Perry et al., 2008; Senathi-Raja et al., 2010; Stavro et al., 2013). Thus, participant performance was unlikely confounded by these factors. Another strength was the use of a urine toxicology screen, which allowed us to examine the effects of METH+ on sustained attention and vigilance, with and without UTOX(meth)+ people.

This study has some limitations. Although we controlled for education, this reflects only one factor of a multifaceted cognitive reserve construct, which buffers against HAND in PWH (Morgan et al., 2012). We did not control for smoking, which has some pro-cognitive effects (Majdi et al., 2019). Although expected, our sample was predominantly male (95%), limiting the generalizability of results to females with HIV. Given the known sex differences in cognitive function among PWH (Heaton et al., 2015; Maki et al., 2018), future research could examine potential three-way interactions between HIV, METH, and sex, within a more representative sample, controlling for cognitive reserve and smoking. Such studies can also be conducted in rodents. Finally, a majority of nadir CD4 values in the present study were self-reported (*n* = 96, 94%, versus *n* = 6, 6% derived from lab values). Given the known memory deficits associated with HIV (Walker & Brown, 2018), the reliability of the nadir CD4 variable in this study is questionable. Future research should obtain information regarding nadir CD4 from more objective sources.

## 5. Conclusions

Controlling for VLS, which was associated with better vigilance, PWH did not exhibit sustained attention and vigilance deficits, but rather, exhibited only psychomotor slowing. Further research should examine whether psychomotor slowing reflects pervasive deficits despite VLS, or an adaptive, speed/ accuracy trade-off. METH+ was associated with deficits in sustained attention and vigilance. This effect was no longer significant after removing recent methamphetamine users, potentially suggesting a sustained attention and vigilance rebound among METH+ people, following abstinence. Further research should examine the direction of this relationship and identify the aspects of recent use (e.g., acute intoxication, residual effects, or withdrawal) that are associated with deficits. These effects can be tested directly in animals. There were no synergistic HIV by METH effects on sustained attention and vigilance. Longitudinal research should examine whether onset of METH+ prior to HIV seroconversion is associated with worse cognitive outcomes. The 5C-CPT was found to be an appropriate measure of sustained attention and vigilance deficits in PWH and METH+ people. Using the 5C-CPT enables future comparisons with rodent research examining the underlying mechanisms driving deficits found in this study. Ultimately, results underscore the need for prompt HIV diagnosis and ART initiation, and treatment of METH+ to preserve cognitive function.

## Funding Sources/Acknowledgments

This study was supported by P50 DA26306 and R01DA043535. Nina Pocuca is also supported by an Interdisciplinary Research Fellowship in NeuroAIDS (IRFN; R25MH081482). We acknowledge Nathan Wood for his contribution to the project.

