## Supplementary material for "Sustained attention and vigilance deficits associated with HIV and a history of methamphetamine dependence": Figures

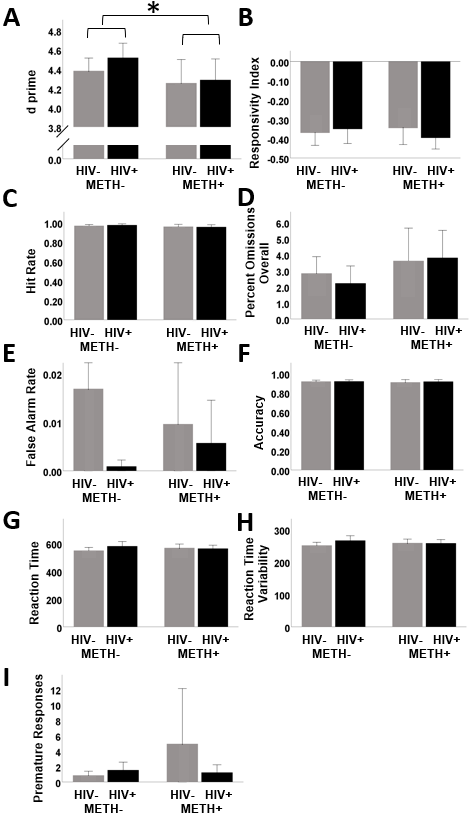

*Figure 1.* Effects of human immunodeficiency virus (HIV) and history of methamphetamine dependence (METH) on 5C-CPT performance.

*N* = 203. METH+ had lower d prime, relative to METH- people (**A**). There were no main or interactive effects of HIV and METH on any of the other outcome variables (**B – I**). Data are presented as means, with error bars representing standard error of the mean. Significant differences are denoted with an asterisk. Corresponding regression estimates, standard errors, and bootstrapped 95% confidence intervals presented in Table 2.

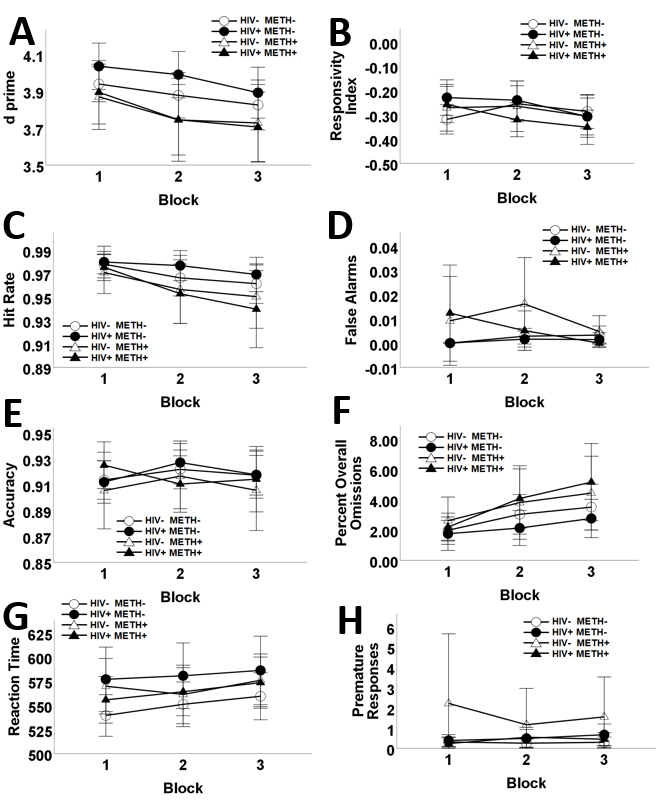

*Figure 2.* Effects of human immunodeficiency virus (HIV) and history of methamphetamine dependence (METH) on 5C-CPT performance across trial blocks.

*N* = 203. Overall performance on the 5C-CPT decreased across blocks. METH+ participants exhibited poorer 5C-CPT performance relative to METH- participants, irrespective of HIV status, as measured by d prime (**A**). METH+ participants had significantly poorer responsivity index, compared to METH- participants, however, this was not significant following Bonferroni correction (**B**). Impaired responding to targets was more exaggerated as time progressed and among METH+ participants, compared to METH- participants (**C**). False alarm rates did not differ across blocks, or according to HIV. METH+ participants had more false alarms compared to METH- participants, however, this effect was not significant follow Bonferroni correction (**D**). HIV+ had lower accuracy, compared to HIV-negative people (**E**). The number of omissions significantly increased across trial blocks. METH+ participants had significantly elevated misses to targets (% omission; **F**). Reaction time slowed over trial blocks and HIV+ had significantly slower reaction time, relative to HIV- subjects (**G**). Finally, premature responses did not differ across blocks, or according to HIV. METH+ participants had significantly more premature responses, however, this effect was not significant following Bonferroni correction (**H**). Data are presented as means, with error bars representing standard error of the mean. Corresponding regression estimates, standard errors, and bootstrapped 95% confidence intervals presented in Table 4.

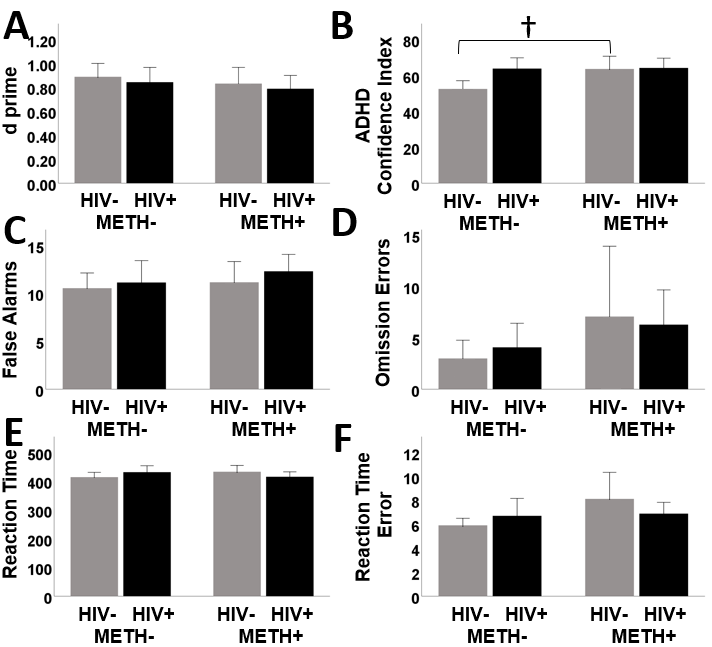

*Figure 3.* Effects of human immunodeficiency virus (HIV) and history of methamphetamine dependence (METH) on Conners’ CPT performance.

*N* = 203. There was a significant HIV x METH interaction on attention-deficit/ hyperactivity disorder (ADHD) confidence index (prior to Bonferroni correction), such that among HIV- people, METH+ was associated with greater ADHD confidence index. METH+ was not associated with ADHD confidence index among people with HIV (**A**). There were no main or interactive effects of HIV and METH on any of the other outcome variables (**A – F**). Data are presented as means, with error bars representing standard error of the mean. Significant difference prior to Bonferroni correction (i.e., *p*<.05) is denoted with a †. Corresponding regression estimates, standard errors, and bootstrapped 95% confidence intervals presented in Table 5.

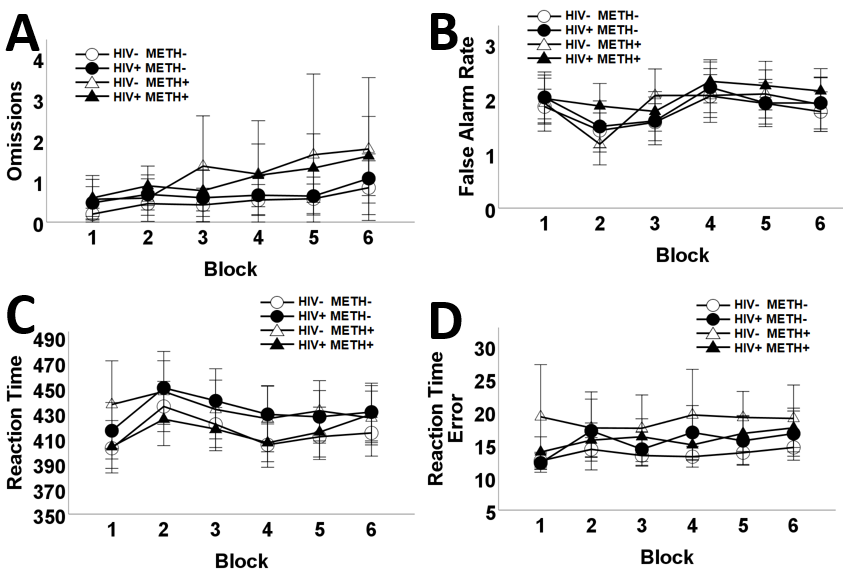

*Figure 4.* Effects of human immunodeficiency virus (HIV) and history of methamphetamine dependence (METH) on Conners’ CPT performance across trial blocks..

*N* = 203. Omissions increased across blocks. METH+ participants exhibited greater omissions, relative to METH- participants (**A**). False alarm rates also increased across blocks; however, false alarm rates did not differ according to HIV or METH status (**B**). HIV+ had significantly slower reaction time compared to HIV- subjects. Similarly, METH+ had significantly slower reaction time compared to METH- participants (**C**). Reaction time error increased across blocks. METH+ participants exhibited greater RT error, than METH- participants (**D**). Data are presented as means, with error bars representing standard error of the mean. Corresponding regression estimates, standard errors, and bootstrapped 95% confidence intervals presented in Table 7.
