## Supplemental Tables for "Sustained attention and vigilance deficits associated with HIV and a history of methamphetamine dependence"

| Supplemental Table 1 | | | | | | | | | |
| --- | --- | --- | --- | --- | --- | --- | --- | --- | --- |
| Main and Interactive Effects of HIV and History of Methamphetamine Dependence on the 5C-CPT (Excluding Participants with a Positive Methamphetamine Urine Toxicology Screen) | | | | | | | | | |
|  | d prime | | | Responsivity Index | | | Hit Rate | | |
| Model 1 | B | SE | 95% CI | B | SE | 95% CI | B | SE | 95% CI |
| Age | -0.083 | 0.047 | [-0.178, 0.006] | -0.018 | 0.019 | [-0.056, 0.021] | -0.008 | 0.005 | [-0.017, 0.001] |
| Gender | **-0.25** | **0.11** | **[-0.462, -0.026]** | -0.027 | 0.055 | [-0.135, 0.086] | -0.001 | 0.01 | [-0.019, 0.019] |
| Education | -0.018 | 0.047 | [-0.106, 0.079] | -0.044 | 0.021 | [-0.085, -0.001]† | 0 | 0.004 | [-0.007, 0.009] |
| VLS | 0.161 | 0.171 | [-0.144, 0.526] | 0.005 | 0.064 | [-0.131, 0.127] | 0.017 | 0.017 | [-0.01, 0.054] |
| HIV | 0.058 | 0.095 | [-0.122, 0.258] | -0.001 | 0.046 | [-0.091, 0.09] | 0.008 | 0.009 | [-0.008, 0.027] |
| METH | -0.055 | 0.083 | [-0.215, 0.107] | -0.028 | 0.04 | [-0.105, 0.048] | -0.001 | 0.007 | [-0.017, 0.011] |
| Model 2 |  |  |  |  |  |  |  |  |  |
| HIVxMETH | -.082 | .160 | [-.380, .241] | -.062 | .073 | [-.212, .076] | -.004 | .015 | [-.032, .027] |
| Model 1 | Percent overall Omissions | | | False Alarm Rate | | | Accuracy | | |
| Age | 0.636 | 0.387 | [-0.128, 1.404] | -0.002 | 0.005 | [-0.013, 0.004] | -0.003 | 0.006 | [-0.014, 0.008] |
| Gender | 0.036 | 0.878 | [-1.834, 1.625] | -0.01 | 0.017 | [-0.052, 0.014] | -0.034 | 0.014 | [-0.063, -0.008]† |
| Education | 0.04 | 0.322 | [-0.627, 0.654] | -0.007 | 0.005 | [-0.018, 0] | 0.006 | 0.005 | [-0.003, 0.018] |
| VLS | -1.471 | 1.444 | [-4.849, 0.899] | -0.002 | 0.004 | [-0.01, 0.005] | -0.007 | 0.013 | [-0.03, 0.021] |
| HIV | -0.723 | 0.846 | [-2.565, 0.8] | -0.015 | 0.015 | [-0.05, 0] | -0.008 | 0.009 | [-0.024, 0.009] |
| METH | 0.153 | 0.661 | [-1.045, 1.547] | -0.008 | 0.012 | [-0.038, 0.007] | 0.005 | 0.009 | [-0.015, 0.022] |
| Model 2 |  |  |  |  |  |  |  |  |  |
| HIVxMETH | .332 | 1.364 | [-2.479, 2.992] | .010 | .016 | [-.012, .047] | .005 | .018 | [-.029, .042] |
| Model 1 | Reaction Time | | | Variability in RT | | | Premature Responses | | |
| Age | 29.632 | 7.285 | [14.603, 43.809] | 12.76 | 3.036 | [6.475, 18.452] | 0.962 | 1.002 | [-0.405, 3.465] |
| Gender | 31.593 | 16.814 | [-1.251, 64.835] | 11.836 | 6.816 | [-1.966, 25.101] | 2.754 | 2.631 | [-0.442, 9.106] |
| Education | 0.056 | 6.567 | [-14.044, 12.935] | 0.393 | 2.792 | [-5.363, 5.894] | -1.442 | 1.47 | [-5.016, 0.251] |
| VLS | 14.563 | 23.303 | [-31.023, 61.175] | 7.974 | 10.317 | [-11.972, 28.444] | 1.103 | 0.76 | [-0.23, 2.988] |
| HIV | 24.709 | 14.99 | [-4.437, 55.799] | 11.868 | 6.605 | [-0.85, 24.64] | -0.071 | 0.793 | [-1.937, 1.222] |
| METH | 6.488 | 13.462 | [-20.364, 32.719] | 3.118 | 6.022 | [-9.359, 14.481] | 1.57 | 1.771 | [-0.705, 5.674] |
| Model 2 |  |  |  |  |  |  |  |  |  |
| HIVxMETH | -29.901 | 26.231 | [-82.582, 20.153] | -12.407 | 11.870 | [-35.652, 10.265] | -3.039 | 2.944 | [-10.015, 0.736] |
| Note. HIV = Human immunodeficiency virus. METH = methamphetamine dependence. VLS = Viral load suppression. Significant effects following bonferroni adjustment (*p=*.006) are bolded. †Significant at *p*<.05 | | | | | | | | | |

| Supplemental Table 2 | | | | |
| --- | --- | --- | --- | --- |
| -2*LL* for the Main Effects and Interaction Models Tested for the 5C-CPT (Excluding Participants with a Positive Methamphetamine Urine Toxicology Screen) | | | | |
|  | Model 1 | Model 2 | χ ^2^ | *p* |
| d prime | 527.088 | 525.871 | 1.217 | .749 |
| Responsivity Index | 59.034 | 53.043 | 5.991 | .112 |
| Hit Rate | -1921.796 | -1922.167 | 0.371 | .946 |
| False Alarm Rate | -2858.879 | -2866.047 | 7.168 | .067 |
| Accuracy | -1807.697 | -1808.201 | 0.504 | .918 |
| Percent overall Omissions | 3079.411 | 3079.049 | 0.362 | .948 |
| Reaction Time | 6099.670 | 6098.213 | 1.457 | .692 |
| Premature Responses | 2507.236 | 2501.855 | 5.381 | .146 |
| Note: Model 1 = Main effects of block, HIV, and history of methamphetamine dependence; Model 2 = 2-way interactions between block, HIV, and history of methamphetamine dependence. Degrees of freedom for model 1 = 10, model 2 = 13. | | | | |

| Supplemental Table 3 | | | | | | | | | | | | | | | |
| --- | --- | --- | --- | --- | --- | --- | --- | --- | --- | --- | --- | --- | --- | --- | --- |
| Main Effects of Block, HIV, and History of Methamphetamine Dependence on the 5C-CPT (Excluding Participants with a Positive Methamphetamine Urine Toxicology Screen) | | | | | | | | | | | | | | | |
|  |  | | d prime | |  |  | | Responsivity Index | |  | |  | Hit Rate | | |
|  | B | SE | | 95% CI | B | | SE | | 95% CI | | B | | | SE | 95% CI |
| Age | **-0.067** | **0.016** | | **[-0.098, -0.036]** | -0.006 | | 0.01 | | [-0.025, 0.014] | | **-0.007** | | | **0.002** | **[-0.011, -0.003]** |
| Gender | **-0.152** | **0.036** | | **[-0.22, -0.081]** | **-0.065** | | **0.026** | | **[-0.113, -0.015]** | | -0.002 | | | 0.004 | [-0.01, 0.006] |
| Education | -0.008 | 0.017 | | [-0.043, 0.027] | **-0.038** | | **0.01** | | **[-0.058, -0.018]** | | 0 | | | 0.002 | [-0.004, 0.003] |
| VLS | **0.14** | **0.052** | | **[0.024, 0.239]** | 0.01 | | 0.032 | | [-0.055, 0.073] | | 0.016 | | | 0.007 | [0, 0.03] |
| Block | **-0.066** | **0.017** | | **[-0.102, -0.03]** | -0.018 | | 0.012 | | [-0.046, 0.008] | | **-0.008** | | | **0.002** | **[-0.012, -0.004]** |
| HIV | 0.046 | 0.03 | | [-0.01, 0.098] | -0.008 | | 0.021 | | [-0.046, 0.03] | | 0.008 | | | 0.004 | [0.001, 0.015] |
| METH | -0.043 | 0.028 | | [-0.097, 0.011] | -0.02 | | 0.02 | | [-0.06, 0.02] | | -0.001 | | | 0.003 | [-0.007, 0.005] |
|  | False Alarms | | | | Accuracy | | | | | | Percent overall Omissions | | | | |
| Age | 0.002 | 0.001 | | [-0.001, 0.003] | -0.003 | | 0.002 | | [-0.006, 0.001] | | **0.636** | | | **0.154** | **[0.288, 0.907]** |
| Gender | 0.006 | 0.002 | | [0, 0.011] | **-0.034** | | **0.005** | | **[-0.044, -0.023]** | | 0.032 | | | 0.336 | [-0.58, 0.696] |
| Education | -0.003 | 0.001 | | [-0.005, -0.001]† | **0.006** | | **0.002** | | **[0.002, 0.011]** | | 0.039 | | | 0.137 | [-0.207, 0.327] |
| VLS | -0.002 | 0.002 | | [-0.005, 0.001] | -0.007 | | 0.005 | | [-0.019, 0.003] | | -1.471 | | | 0.638 | [-2.743, -0.085]† |
| Block | 0 | 0.001 | | [-0.002, 0.002] | 0.002 | | 0.002 | | [-0.002, 0.006] | | **0.726** | | | **0.172** | **[0.398, 1.076]** |
| HIV | -0.001 | 0.001 | | [-0.003, 0.001] | -0.008 | | 0.004 | | [-0.015, 0.000] | | -0.733 | | | 0.304 | [-1.284, -0.099]† |
| METH | 0.003 | 0.002 | | [-0.002, 0.006] | 0.005 | | 0.004 | | [-0.001, 0.013] | | 0.158 | | | 0.252 | [-0.34, 0.631] |
|  | Reaction Time | | | | Premature Responses | | | | | |  | | | | |
| Age | **29.63** | **2.64** | | **[24.191, 34.892]** | 0.321 | | 0.115 | | [-0.032, 0.546] | |  | | |  |  |
| Gender | **31.483** | **5.512** | | **[19.746, 42.658]** | 0.918 | | 0.277 | | [0.047, 1.456]† | |  | | |  |  |
| Education | 0.073 | 2.303 | | [-5.044, 4.738] | -0.481 | | 0.154 | | [-0.748, -0.017]† | |  | | |  |  |
| VLS | 15.209 | 7.248 | | [-0.098, 30.253] | 0.368 | | 0.09 | | [0.176, 0.552]† | |  | | |  |  |
| Block | 6.837 | 2.438 | | [1.795, 11.912]† | -0.032 | | 0.092 | | [-0.242, 0.154] | |  | | |  |  |
| HIV | **24.561** | **4.868** | | **[14.395, 34.026]** | -0.024 | | 0.097 | | [-0.197, 0.164] | |  | | |  |  |
| METH | 6.74 | 4.41 | | [-1.254, 15.177] | 0.523 | | 0.189 | | [-0.037, 0.879] | |  | | |  |  |
| Note. HIV = Human immunodeficiency virus. METH = methamphetamine dependence. VLS = Viral load suppression. Significant effects following bonferroni adjustment (*p=*.006) are bolded. †Significant at *p*=.05 | | | | | | | | | | | | | | | |

| Supplemental Table 4 | | | | | | | | | | | | | | |
| --- | --- | --- | --- | --- | --- | --- | --- | --- | --- | --- | --- | --- | --- | --- |
| Main and Interactive Effects of HIV and History of Methamphetamine Dependence on Conners’ CPT (Excluding Participants with a Positive Methamphetamine Urine Toxicology Screen) | | | | | | | | | | | | | | |
|  |  | | d prime | |  |  | | ADHD Confidence Index | |  |  | | False Alarms | |
| Model 1 | B | SE | | 95% CI | B | | SE | | 95% CI | B | | SE | | 95% CI |
| Age | 0.057 | 0.03 | | [-0.003, 0.113] | **6.757** | | **1.42** | | **[3.848, 9.595]** | **-1.339** | | **0.469** | | **[-2.286, -0.492]** |
| Gender | 0.148 | 0.081 | | [-0.001, 0.317] | **-13.239** | | **3.485** | | **[-19.671, -6.207]** | -2.719 | | 1.017 | | [-4.713, -0.648]† |
| Education | 0.044 | 0.032 | | [-0.018, 0.106] | 0.468 | | 1.512 | | [-2.724, 3.381] | -0.777 | | 0.511 | | [-1.724, 0.25] |
| VLS | -0.007 | 0.123 | | [-0.252, 0.21] | 0.05 | | 5.02 | | [-10.738, 9.48] | -0.011 | | 1.74 | | [-3.437, 3.63] |
| HIV | -0.02 | 0.071 | | [-0.159, 0.119] | 1.861 | | 2.981 | | [-4.273, 7.643] | 0.393 | | 1.087 | | [-1.801, 2.563] |
| METH | 0.027 | 0.062 | | [-0.106, 0.146] | 4.872 | | 2.79 | | [-0.453, 10.689] | -0.859 | | 1.014 | | [-2.826, 1.152] |
| Model 2 |  |  | |  |  | |  | |  |  | |  | |  |
| HIVxMETH | -.012 | .123 | | [-.252, .234] | -12.198 | | 5.442 | | [-22.508, -1.636]† | .626 | | 1.886 | | [-2.861, 4.384] |
| Model 1 | Omission Errors | | | | Hit Reaction Time | | | | | Reaction Time Error | | | | |
| Age | 0.986 | 0.676 | | [-0.318, 2.328] | **25.306** | | **4.393** | | **[16.886, 33.719]** | 0.435 | | 0.202 | | [0.018, 0.838]† |
| Gender | -3.016 | 2.028 | | [-7.468, 0.366] | 15.001 | | 12.277 | | [-11.088, 39.483] | -0.084 | | 0.517 | | [-1.038, 0.983] |
| Education | -1.32 | 0.951 | | [-3.467, 0.233] | -2.274 | | 5.027 | | [-12.945, 7.107] | 0.003 | | 0.319 | | [-0.589, 0.704] |
| VLS | -2.421 | 2.545 | | [-7.578, 2.143] | 11.389 | | 14.409 | | [-14.947, 40.167] | 0.52 | | 0.66 | | [-0.757, 1.895] |
| HIV | -1.204 | 2.002 | | [-5.531, 2.144] | 7.467 | | 10.81 | | [-12.703, 29.154] | 0.271 | | 0.616 | | [-0.797, 1.649] |
| METH | 0.247 | 1.654 | | [-2.539, 3.814] | 2.108 | | 9.923 | | [-16.855, 22.229] | 0.246 | | 0.458 | | [-0.658, 1.148] |
| Model 2 |  |  | |  |  | |  | |  |  | |  | |  |
| HIVxMETH | -2.749 | 3.837 | | [-11.320, 3.466] | -25.282 | | 18.265 | | [-60.502, 10.572] | -1.397 | | 1.138 | | [-3.923, .512] |
| Note. HIV = Human immunodeficiency virus. METH = methamphetamine dependence. VLS = Viral load suppression. Significant effects following bonferroni adjustment (*p=*.006) are bolded. †Significant at *p*=.05 | | | | | | | | | | | | | | |

| Supplemental Table 5 | | | | |
| --- | --- | --- | --- | --- |
| -2*LL* for the Main Effects and Interaction Models Tested for Conners’ CPT (Excluding Participants with a Positive Methamphetamine Urine Toxicology Screen) | | | | |
|  | Model 1 | Model 2 | χ^2^ | *p* |
| Omissions | 4397.344 | 4392.501 | 4.843 | .184 |
| False Alarm Rate | 3572.464 | 3571.263 | 1.201 | .753 |
| Reaction Time | 11594.240 | 11588.337 | 5.903 | .116 |
| Reaction Time Error | 7504.518 | 7499.845 | 4.673 | .197 |
| Note: Model 1 = Main effects of block, HIV, and history of methamphetamine dependence; Model 2 = 2-way interactions between block, HIV, and history of methamphetamine dependence. Degrees of freedom for model 1 = 10, model 2 = 13. | | | | |

| Supplemental Table 6 | | | | | | | | | | | | | | | | | | |
| --- | --- | --- | --- | --- | --- | --- | --- | --- | --- | --- | --- | --- | --- | --- | --- | --- | --- | --- |
| Main Effects of Block, HIV, and History of Methamphetamine Dependence on Conners’ CPT (Excluding Participants with a Positive Methamphetamine Urine Toxicology Screen) | | | | | | | | | | | | | | | | | | |
|  |  | Omissions | | | | |  | |  | False Alarm Rate | | | |  |  | Reaction Time | | |
|  | B | | SE | 95% CI | B | | | | | | SE | 95% CI | B | | | | SE | 95% CI |
| Age | **0.164** | | **0.046** | **[0.073, 0.257]** | **-0.223** | | | | | | **0.034** | **[-0.296, -0.154]** | **25.393** | | | | **1.031** | **[23.344, 27.52]** |
| Gender | **-0.503** | | **0.118** | **[-0.71, -0.266]** | **-0.453** | | | | | | **0.079** | **[-0.606, -0.299]** | **15.347** | | | | **2.74** | **[9.666, 20.803]** |
| Education | **-0.22** | | **0.047** | **[-0.3, -0.127]** | **-0.129** | | | | | | **0.035** | **[-0.193, -0.066]** | -2.127 | | | | 1.15 | [-4.417, -0.046]† |
| VLS | -0.404 | | 0.161 | [-0.731, -0.122]† | -0.002 | | | | | | 0.107 | [-0.209, 0.197] | **11.226** | | | | **4.186** | **[2.357, 19.238]** |
| Block | **0.118** | | **0.031** | **[0.062, 0.179]** | 0.052 | | | | | | 0.018 | [0.015, 0.089]† | -0.04 | | | | 0.674 | [-1.521, 1.496] |
| HIV | -0.201 | | 0.108 | [-0.411, 0.024] | 0.066 | | | | | | 0.073 | [-0.073, 0.201] | **7.712** | | | | **2.407** | **[3.036, 12.315]** |
| METH | 0.041 | | 0.093 | [-0.152, 0.228] | -0.143 | | | | | | 0.068 | [-0.273, 0.015] | 1.721 | | | | 2.195 | [-2.958, 6.26] |
|  |  | Reaction Time Error | | | |  | |  | | |  |  |  | | | |  |  |
| Age | **1.069** | | **0.163** | **[0.782, 1.377]** |  | | | | | |  |  |  | | | |  |  |
| Gender | -0.195 | | 0.364 | [-0.895, 0.514] |  | | | | | |  |  |  | | | |  |  |
| Education | -0.033 | | 0.225 | [-0.576, 0.441] |  | | | | | |  |  |  | | | |  |  |
| VLS | 1.492 | | 0.577 | [0.222, 2.618]† |  | | | | | |  |  |  | | | |  |  |
| Block | **0.416** | | **0.105** | **[0.219, 0.6]** |  | | | | | |  |  |  | | | |  |  |
| HIV | 0.68 | | 0.425 | [-0.22, 1.473] |  | | | | | |  |  |  | | | |  |  |
| METH | 0.509 | | 0.329 | [-0.095, 1.133] |  | | | | | |  |  |  | | | |  |  |
| Note. HIV = Human immunodeficiency virus. METH = methamphetamine dependence. VLS = Viral load suppression. Significant effects following bonferroni adjustment (*p=*.006) are bolded. †Significant at *p*=.05 | | | | | | | | | | | | | | | | | | |
