## Supplementary material for "Sustained attention and vigilance deficits associated with HIV and a history of methamphetamine dependence": Tables

| Table 1 | | | | | |
| --- | --- | --- | --- | --- | --- |
| Sample characteristics | | | | | |
|  | HIV-/METH-  (n = 67; A) | HIV+/METH-  (n = 49; B) | HIV-/METH+  (n = 36; C) | HIV+/METH+  (n = 53; D) | Group Differences |
| Age | 44.60 (16.27) | 47.27 (14.70) | 43.28 (12.26) | 40.85 (8.45) | A B > C D |
| Gender (% males; ref) | 41 (61%) | 45 (92%) | 20 (56%) | 52 (98%) | A C < B D |
| Education | 14.69 (1.80) | 14.61(2.33) | 12.75(3.07) | 13.98 (2.27) | A B > C D |
| Ethnicity |  |  |  |  |  |
| % White (ref) | 37 (55%) | 27 (55%) | 19 (53%) | 31 (59%) | - |
| % Black | 8 (12%) | 9 (18%) | 6 (17%) | 5 (9%) | - |
| % Hispanic | 18 (27%) | 9 (18%) | 8 (22%) | 15 (28%) | - |
| % Other^a^ | 4 (6%) | 4 (8%) | 3 (8%) | 2 (4%) | - |
| Mental Health^b^ |  |  |  |  |  |
| ASPD | 3 (5%) | 2 (4%) | 11 (31%) | 9 (17%) | A B < C D |
| ADHD (Current) | 1 (2%) | 0 | 1 (3%) | 1 (2%) | - |
| ADHD(Lifetime) | 1 (2%) | 3 (6%) | 4 (11%) | 3 (6%) | - |
| Current nicotine dependence | 2 (3%) | 0 | 3 (8%) | 6 (11%) | A B < C D |
| Lifetime dependence |  |  |  |  |  |
| Alcohol | 3 (5%) | 6 (12%) | 12 (33%) | 12 (23%) | A B < C D |
| Cannabis | 2 (3%) | 2 (4%) | 3 (8%) | 7 (13%) | - |
| Cocaine | 1 (2%) | 3 (6%) | 6 (17%) | 7 (13%) | A B < C D |
| Opioid | 1 (2%) | 0 | 5 (14%) | 2 (4%) | A B < C D |
| Positive urine toxicology |  |  |  |  |  |
| Meth | 0 | 0 | 5 (14%) | 11 (21%) | - |
| THC | 2 (3%) | 7 (14%) | 2 (6%) | 5 (9%) | B D > A C |
| Amphetamine | 0 | 1 (2%) | 4 (11%) | 11 (20%) | A B < C D |
| Opiates | 0 | 5 (10%) | 0 | 0 | B > A C D |
| Benzodiazepines | 0 | 2 (4%) | 2 (6%) | 3 (6%) | - |
| Methamphetamine Characteristics |  |  |  |  |  |
| Current METH+ | - | - | 7 (19%) | 8 (15%) | - |
| Ever used Meth | 2 | 6 | 36 | 53 | A B < C D |
| Age at first use | 32.50 (3.54) | 28.67 (7.29) | 26.08 (13.23) | 24.92 (7.89) | - |
| Days since use | 5387.29 (7360.52) | 1327.04 (1966.95) | 162.97 (366.73) | 139.40 (160.67) | A B > C D |
| Cumulative  duration (days) | 176.00 (120.21) | 182.80 (188.30) | 2980.64 (2753.166) | 2073.61 (1885.19) | A B < C D |
| Cumulative  quantity (grams) | 916.29 (1286.81) | 75.02 (141.99) | 4568.91 (8041.65) | 2668.86 (4406.01) | A B < C D |
| Current IV use | - | - | 3 (8.3%) | 7 (13%) | - |
| HIV characteristics |  |  |  |  |  |
| Currently on ART | - | 44 (90%) | - | 46 (87%) | - |
| Virally suppressed | - | 43 (88%) | - | 35 (66%) | B > D |
| Nadir CD4 | - | 319.84 (220.809) | - | 311.60 (190.539) | - |
| *Note*. Unless otherwise stated, binary variables are coded 0 = no, 1 = yes. HIV = Human Immunodeficiency Virus. METH = History of methamphetamine dependence. ASPD = Antisocial personality disorder. ADHD = Attention-deficit/ hyperactivity disorder. THC = THC = Δ-9 Tetrahydrocannabinol. IV = intravenous.  ^a^Asian merged with “other” for multinominal logistic regression due to sparsity of category  ^b^Assessed using the Diagnostic Interview Schedule for DSM-IV | | | | | |

| Table 2 | | | | | | | | | |
| --- | --- | --- | --- | --- | --- | --- | --- | --- | --- |
| Main and Interactive Effects of HIV and History of Methamphetamine Dependence on the 5C-CPT | | | | | | | | | |
|  | d prime | | | Responsivity Index | | | Hit Rate | | |
| Model 1 | B | SE | 95% CI | B | SE | 95% CI | B | SE | 95% CI |
| Age | -0.081 | 0.048 | [-0.173, 0.017] | -0.02 | 0.018 | [-0.055, 0.017] | -0.007 | 0.005 | [-0.016, 0.002] |
| Gender | **-0.315** | **0.111** | **[-0.556, -0.109]** | -0.029 | 0.05 | [-0.128, 0.07] | -0.008 | 0.01 | [-0.027, 0.01] |
| Education | -0.051 | 0.061 | [-0.173, 0.07] | -0.036 | 0.018 | [-0.073, 0.001] | -0.004 | 0.006 | [-0.017, 0.007] |
| VLS | 0.151 | 0.16 | [-0.165, 0.458] | 0.028 | 0.054 | [-0.082, 0.129] | 0.014 | 0.017 | [-0.017, 0.052] |
| HIV | 0.023 | 0.095 | [-0.169, 0.202] | -0.007 | 0.042 | [-0.09, 0.076] | 0.003 | 0.01 | [-0.016, 0.024] |
| METH | **-0.219** | **0.1** | **[-0.43, -0.025]** | -0.034 | 0.038 | [-0.107, 0.041] | -0.018 | 0.01 | [-0.04, 0.002] |
| Model 2 |  |  |  |  |  |  |  |  |  |
| HIVxMETH | -.125 | .166 | [-.456, .181] | -.061 | .069 | [-.199, .070] | -.010 | .017 | [-.046, .024] |
| Model 1 | Percent Omissions Overall | | | False Alarm Rate | | | Accuracy | | |
| Age | 0.625 | 0.38 | [-0.148, 1.383] | -0.003 | 0.004 | [-0.012, 0.004] | -0.003 | 0.005 | [-0.013, 0.007] |
| Gender | 0.623 | 0.827 | [-1.038, 2.215] | -0.008 | 0.015 | [-0.044, 0.014] | -0.037 | 0.012 | [-0.062, -0.013]† |
| Education | 0.31 | 0.464 | [-0.554, 1.281] | -0.003 | 0.005 | [-0.013, 0.006] | 0.002 | 0.007 | [-0.012, 0.014] |
| VLS | -1.381 | 1.427 | [-4.492, 1.138] | 0.003 | 0.005 | [-0.004, 0.014] | -0.011 | 0.012 | [-0.035, 0.013] |
| HIV | -0.407 | 0.858 | [-2.202, 1.194] | -0.012 | 0.013 | [-0.044, 0.005] | -0.012 | 0.009 | [-0.032, 0.005] |
| METH | 1.417 | 0.786 | [-0.103, 3.021] | -0.003 | 0.011 | [-0.029, 0.013] | -0.007 | 0.011 | [-0.028, 0.014] |
| Model 2 |  |  |  |  |  |  |  |  |  |
| HIVxMETH | .751 | 1.488 | [-2.561, 3.354] | .013 | .015 | [-.011, .047] | -.003 | .018 | [-.039, .034] |
| Model 1 | Reaction Time | | | Variability in RT | | | Premature Responses | | |
| Age | **32.348** | **6.821** | **[18.367, 45.442]** | **14.058** | **2.919** | **[8.079, 19.78]** | 0.913 | 0.862 | [-0.323, 2.915] |
| Gender | 27.172 | 16.094 | [-4.503, 59.139] | 9.38 | 7.205 | [-4.522, 23.772] | 2.678 | 2.121 | [-0.247, 7.428] |
| Education | -5.017 | 6.094 | [-17.027, 6.569] | -2.548 | 2.921 | [-8.681, 3.107] | -1.229 | 1.225 | [-4.124, 0.414] |
| VLS | -6.574 | 25.107 | [-55.877, 40.452] | -2.188 | 10.849 | [-24.827, 17.636] | 1.613 | 0.709 | [0.293, 3.052]† |
| HIV | 24.917 | 15.06 | [-5.39, 54.275] | 11.557 | 6.499 | [-0.807, 24.223] | 0.328 | 0.783 | [-1.542, 1.665] |
| METH | 7.168 | 12.91 | [-18.252, 33.525] | 2.173 | 5.957 | [-9.214, 14.164] | 1.623 | 1.398 | [-0.528, 4.927] |
| Model 2 |  |  |  |  |  |  |  |  |  |
| HIVxMETH | -19.096 | 26.171 | [-71.094, 31.514] | -8.317 | 11.634 | [-31.052, 13.541] | -2.892 | 2.642 | [-9.162, .685] |
| Note. HIV = human immunodeficiency virus. METH = methamphetamine dependence. VLS = Viral load suppression. Significant effects following bonferroni adjustment (*p=*.006) are bolded. †Significant at *p*<.05 | | | | | | | | | |

| Table 3 | | | | |
| --- | --- | --- | --- | --- |
| -2*LL* for the Main Effects and Interaction Models Tested for the 5C-CPT | | | | |
|  | Model 1 | Model 2 | χ^2^ | *p* |
| d prime | 652.333 | 650.851 | 1.482 | .686 |
| Responsivity Index | 64.725 | 58.413 | 6.312 | .097 |
| Hit Rate | -1884.082 | -1889.028 | 4.946 | .176 |
| False Alarm Rate | -2626.580 | -2633.939 | 7.359 | .061 |
| Accuracy | -1912.428 | -1915.171 | 2.743 | .433 |
| Percent overall Omissions | 3515.696 | 3511.669 | 4.027 | .259 |
| Reaction Time | 6675.166 | 6674.505 | 0.661 | .882 |
| Premature Responses | 2735.521 | 2730.309 | 5.212 | .157 |
| Note: Model 1 = Main effects of block, HIV, and history of methamphetamine dependence; Model 2 = 2-way interactions between block, HIV, and history of methamphetamine dependence. Degrees of freedom for model 1 = 10, model 2 = 13. | | | | |

| Table 4 | | | | | | | | | | | | | | |
| --- | --- | --- | --- | --- | --- | --- | --- | --- | --- | --- | --- | --- | --- | --- |
| Main Effects of Block, HIV, and History of Methamphetamine Dependence on the 5C-CPT | | | | | | | | | | | | | | |
|  |  | d prime | | |  |  | Responsivity Index | | |  |  | Hit Rate | | |
|  | B | | SE | 95% CI | | B | | SE | 95% CI | | B | | SE | 95% CI |
| Age | **-0.069** | | **0.016** | **[-0.099, -0.039]** | | -0.009 | | 0.01 | [-0.028, 0.01] | | **-0.007** | | **0.002** | **[-0.011, -0.003]** |
| Gender | **-0.204** | | **0.036** | **[-0.274, -0.129]** | | **-0.064** | | **0.025** | **[-0.11, -0.016]** | | -0.009 | | 0.004 | [-0.017, -0.001]† |
| Education | -0.033 | | 0.018 | [-0.065, 0.003] | | **-0.031** | | **0.009** | **[-0.05, -0.012]** | | -0.004 | | 0.003 | [-0.009, 0.001] |
| VLS | **0.136** | | **0.05** | **[0.027, 0.240]** | | 0.039 | | 0.028 | [-0.015, 0.093] | | 0.014 | | 0.007 | [0.001, 0.027]† |
| Block | **-0.072** | | **0.017** | **[-0.106, -0.037]** | | -0.019 | | 0.012 | [-0.041, 0.002] | | **-0.01** | | **0.002** | **[-0.015, -0.006]** |
| HIV | 0.016 | | 0.032 | [-0.047, 0.079] | | -0.005 | | 0.02 | [-0.042, 0.029] | | 0.003 | | 0.004 | [-0.004, 0.011] |
| METH | **-0.173** | | **0.03** | **[-0.228, -0.112]** | | -0.039 | | 0.019 | [-0.076, -0.001]† | | **-0.018** | | **0.004** | **[-0.025, -0.009]** |
|  | False Alarms | | | | | Accuracy | | | | | Percent overall Omissions | | | |
| Age | 0.001 | | 0.001 | [-0.001, 0.002] | | -0.003 | | 0.002 | [-0.007, 0.001] | | **0.626** | | **0.159** | **[0.317, 0.924]** |
| Gender | 0.007 | | 0.002 | [0.002, 0.011]† | | **-0.037** | | **0.005** | **[-0.046, -0.028]** | | 0.615 | | 0.373 | [-0.107, 1.379] |
| Education | 0 | | 0.002 | [-0.004, 0.004] | | 0.002 | | 0.002 | [-0.003, 0.007] | | 0.31 | | 0.199 | [-0.124, 0.671] |
| VLS | 0.003 | | 0.003 | [-0.002, 0.009] | | -0.011 | | 0.006 | [-0.022, 0.001] | | -1.385 | | 0.616 | [-2.583, -0.088]† |
| Block | -0.001 | | 0.001 | [-0.005, 0.002] | | 0 | | 0.002 | [-0.004, 0.004] | | **0.907** | | **0.19** | **[0.568, 1.281]** |
| HIV | 0.001 | | 0.002 | [-0.003, 0.005] | | **-0.012** | | **0.004** | **[-0.018, -0.005]** | | -0.416 | | 0.345 | [-1.016, 0.175] |
| METH | 0.007 | | 0.003 | [0.001, 0.013]† | | -0.007 | | 0.004 | [-0.014, 0.001] | | **1.422** | | **0.327** | **[0.688, 2.053]** |
|  | Reaction Time | | | | | Premature Responses | | | | |  | | | |
| Age | **32.351** | | **2.544** | **[26.975, 37.558]** | | 0.304 | | 0.108 | [-0.017, 0.52] | |  | |  |  |
| Gender | **27.126** | | **5.125** | **[16.443, 37.596]** | | 0.893 | | 0.246 | [0.112, 1.369]† | |  | |  |  |
| Education | -4.909 | | 2.067 | [-9.134, -0.432]† | | -0.41 | | 0.143 | [-0.625, -0.096]† | |  | |  |  |
| VLS | -6.14 | | 7.7 | [-19.527, 9.748] | | **0.538** | | **0.099** | **[0.315, 0.726]** | |  | |  |  |
| Block | **7.191** | | **2.318** | **[2.351, 11.694]** | | 0 | | 0.091 | [-0.188, 0.168] | |  | |  |  |
| HIV | **24.765** | | **4.891** | **[15.298, 34.334]** | | 0.109 | | 0.109 | [-0.097, 0.313] | |  | |  |  |
| METH | 7.505 | | 4.242 | [-1.243, 16.52] | | 0.541 | | 0.171 | [0.047, 0.868]† | |  | |  |  |
| Note. HIV = human immunodeficiency virus. METH = methamphetamine dependence. VLS = Viral load suppression. Significant effects following bonferroni adjustment (*p=*.006) are bolded. †Significant at *p*=.05 | | | | | | | | | | | | | | |

| Table 5 | | | | | | | | | | | | | | | |
| --- | --- | --- | --- | --- | --- | --- | --- | --- | --- | --- | --- | --- | --- | --- | --- |
| Main and Interactive Effects of HIV and History of Methamphetamine Dependence on Conners’ CPT | | | | | | | | | | | | | | | |
|  |  | d prime | | |  |  | ADHD Confidence Index | | | |  |  | False Alarms | | |
| Model 1 | B | | SE | 95% CI | B | | | SE | 95% CI | B | | | | SE | 95% CI |
| Age | 0.055 | | 0.03 | [-0.006, 0.114] | **6.871** | | | **1.339** | **[4.274, 9.35]** | **-1.245** | | | | **0.495** | **[-2.267, -0.316]** |
| Gender | 0.112 | | 0.076 | [-0.043, 0.255] | **-13.126** | | | **3.396** | **[-20.108, -6.178]** | -2.19 | | | | 1.124 | [-4.382, 0.089] |
| Education | 0.02 | | 0.031 | [-0.044, 0.079] | 0.928 | | | 1.411 | [-2.037, 3.487] | -0.323 | | | | 0.535 | [-1.282, 0.812] |
| VLS | -0.026 | | 0.102 | [-0.242, 0.164] | 0.092 | | | 4.675 | [-9.946, 8.915] | 0.68 | | | | 1.524 | [-2.425, 3.552] |
| HIV | -0.015 | | 0.07 | [-0.147, 0.126] | 1.741 | | | 2.988 | [-4.526, 7.371] | 0.276 | | | | 1.128 | [-2.039, 2.398] |
| METH | -0.03 | | 0.062 | [-0.147, 0.091] | 8.133 | | | 2.712 | [2.878, 13.529]† | 0.404 | | | | 1.074 | [-1.654, 2.524] |
| Model 2 |  | |  |  |  | | |  |  |  | | | |  |  |
| HIVxMETH | .022 | | .123 | [-.224, .259] | -10.610 | | | 5.286 | [-21.542, -1.050]† | .098 | | | | 2.001 | [-3.802, 4.083] |
| Model 1 | Omission Errors | | | | Reaction Time | | | | | Reaction Time Error | | | | | |
| Age | 0.892 | | 0.738 | [-0.515, 2.444] | **25.215** | | | **4.351** | **[16.68, 33.62]** | 0.473 | | | | 0.232 | [0.003, 0.903]† |
| Gender | -2.398 | | 2.17 | [-7.168, 1.152] | 12.836 | | | 11.829 | [-11.074, 37.625] | -0.325 | | | | 0.693 | [-1.785, 0.916] |
| Education | -0.448 | | 1.086 | [-2.614, 1.589] | -3.702 | | | 4.965 | [-14.067, 5.185] | -0.038 | | | | 0.315 | [-0.613, 0.63] |
| VLS | -3.255 | | 3.102 | [-9.429, 2.421] | 3.105 | | | 15.591 | [-26.314, 33.667] | 0.033 | | | | 0.987 | [-1.951, 1.876] |
| HIV | -1.164 | | 2.263 | [-6.118, 2.595] | 7.919 | | | 10.467 | [-12.425, 29.801] | -0.184 | | | | 0.767 | [-1.741, 1.263] |
| METH | 2.827 | | 1.971 | [-0.569, 7.193] | 7.087 | | | 9.928 | [-12.372, 26.677] | 1.293 | | | | 0.632 | [0.127, 2.57]† |
| Model 2 |  | |  |  |  | | |  |  |  | | | |  |  |
| HIVxMETH | -2.304 | | 3.952 | [-10.777, 4.530] | -21.667 | | | 19.296 | [-59.462, 18.401] | -1.962 | | | | 1.501 | [-5.218, .760] |
| Note. HIV = Human immunodeficiency virus. METH = methamphetamine dependence. VLS = Viral load suppression. Significant effects following bonferroni adjustment (*p=*.006) are bolded. †Significant at *p*=.05 | | | | | | | | | | | | | | | |

| Table 6 | | | | |
| --- | --- | --- | --- | --- |
| -2*LL* for the Main Effects and Interaction Models Tested for Conners’ CPT | | | | |
|  | Model 1 | Model 2 | χ^2^ | *p* |
| Omissions | 4976.467 | 4969.703 | 6.764 | .080 |
| False Alarm Rate | 3926.818 | 3926.126 | 0.692 | .875 |
| Reaction Time | 12839.189 | 12832.874 | 6.315 | .097 |
| Reaction Time Error | 8555.174 | 8549.930 | 5.244 | .155 |
| Note: Model 1 = Main effects of block, HIV, and history of methamphetamine dependence; Model 2 = 2-way interactions between block, HIV, and history of methamphetamine dependence. Degrees of freedom for model 1 = 10, model 2 = 13. | | | | |

| Table 7 | | | | | | | | | | | | | | | |
| --- | --- | --- | --- | --- | --- | --- | --- | --- | --- | --- | --- | --- | --- | --- | --- |
| Main Effects of Block, HIV, and History of Methamphetamine Dependence on Conners’ CPT | | | | | | | | | | | | | | | |
|  |  | Omissions | | | |  |  | False Alarm Rate | | |  |  | Reaction Time | | |
|  | B | | SE | 95% CI | | | B | | SE | 95% CI | | B | | SE | 95% CI |
| Age | **0.149** | | **0.049** | **[0.063, 0.246]** | | | **-0.207** | | **0.032** | **[-0.268, -0.141]** | | **25.258** | | **1.076** | **[23.332, 27.225]** |
| Gender | **-0.4** | | **0.122** | **[-0.621, -0.179]** | | | **-0.365** | | **0.08** | **[-0.525, -0.206]** | | **13.025** | | **2.713** | **[7.449, 18.507]** |
| Education | -0.075 | | 0.052 | [-0.17, 0.024] | | | -0.054 | | 0.033 | [-0.12, 0.012] | | **-3.5** | | **1.164** | **[-5.62, -1.294]** |
| VLS | **-0.543** | | **0.186** | **[-0.903, -0.174]** | | | 0.113 | | 0.101 | [-0.086, 0.296] | | 3.021 | | 4.023 | [-4.895, 10.836] |
| Block | **0.149** | | **0.033** | **[0.091, 0.208]** | | | **0.054** | | **0.017** | **[0.019, 0.088]** | | -0.208 | | 0.732 | [-1.661, 1.262] |
| HIV | -0.194 | | 0.104 | [-0.391, 0.011] | | | 0.046 | | 0.069 | [-0.1, 0.193] | | **7.995** | | **2.568** | **[3.224, 13.012]** |
| METH | **0.471** | | **0.097** | **[0.271, 0.669]** | | | 0.067 | | 0.064 | [-0.06, 0.199] | | **7.053** | | **2.339** | **[2.348, 11.677]** |
|  |  | Reaction Time Error | | |  | |  | |  |  | |  | |  |  |
| Age | **1.15** | | **0.167** | **[0.822, 1.451]** | | |  | |  |  | |  | |  |  |
| Gender | -0.821 | | 0.476 | [-1.687, 0.146] | | |  | |  |  | |  | |  |  |
| Education | -0.125 | | 0.21 | [-0.543, 0.272] | | |  | |  |  | |  | |  |  |
| VLS | 0.199 | | 0.611 | [-1.058, 1.466] | | |  | |  |  | |  | |  |  |
| Block | **0.394** | | **0.124** | **[0.162, 0.623]** | | |  | |  |  | |  | |  |  |
| HIV | -0.397 | | 0.456 | [-1.339, 0.58] | | |  | |  |  | |  | |  |  |
| METH | **2.939** | | **0.402** | **[2.117, 3.731]** | | |  | |  |  | |  | |  |  |
| Note. HIV = human immunodeficiency virus. METH = methamphetamine dependence. VLS = Viral load suppression. Significant effects following bonferroni adjustment (*p=*.006) are bolded. †Significant at *p*=.05 | | | | | | | | | | | | | | | |
